# Neural competitive queuing of ordinal structure underlies skilled sequential action

**DOI:** 10.1101/383364

**Authors:** Katja Kornysheva, Dan Bush, Sophie S Meyer, Anna Sadnicka, Gareth Barnes, Neil Burgess

**Author notes:** **Corresponding author**: Dr Katja Kornysheva, Lecturer (Assistant Professor), School of Psychology, Bangor University, Wales LL57 2AS, United Kingdom.

## Abstract

The fluent retrieval and production of movement sequences is essential for a variety of daily activities such as speech, tool-use, musical and athletic performance, but the neural mechanisms underlying sequence planning remain elusive. Here, participants learned sequences of finger presses with different timings and different finger orders, and reproduced them in a magneto-encephalography (MEG) scanner. We classified the MEG patterns immediately preceding each press in the sequence, and examined their dynamics over the production of the whole sequence. Our results confirm a role for the ‘competitive queuing’ of upcoming action representations in the production of learned motor sequences, extending previous computational and non-human primate recording studies to non-invasive measures in humans. In addition, we show that competitive queuing does not simply reflect specific motor actions, but representations of higher-level sequential order that generalise across different motor sequences. Finally, we show that the quality of competitive queuing predicts participants’ production accuracy, and originates from parahippocampal and cerebellar sources. These results suggest that the brain learns and produces multiple behavioural sequences by flexibly combining representations of specific actions with more abstract, parallel representations of sequential structure.

## Main

The majority of skilled human behaviours such as speech, handwriting, playing a musical instrument and athletic performance evolve in temporally structured sequences. A breakdown of their fluency e.g. in dyspraxia, stuttering and task-dependent dystonia can lead to a profound impairment of the individual’s everyday functioning^1-5^. Historically two opposing theories have been proposed for the neural basis of motor sequence control. The associative chaining account originating from the pioneering work of Ebbinghaus^6^ in 1885 proposed strong forward connections between successive elements of a sequence, leading to the behaviorist idea that each sequence element serves as a conditioning stimulus for the subsequent element during the formation of complex behaviours^7^. In modern neuroscience this hypothesis has engendered the formulation of state-space models of spatio-temporal motor control, including of skilled movement sequences such as writing and Morse code production.^8-11^ These models are characterized by a spatio-temporal trajectory that is determined by the serial evolution of population activity, such that a population state *n* triggers the state *n+1*, the latter *n+2* and so forth. In recurrent neural networks (RNN) inspired by the neocortex, the evolution of multiunit activity is primarily determined by the connectivity matrix as acquired through learning. In other words, spatio-temporal movements are controlled by a serial transition through neural population states (multi-unit trajectory), which are mapped onto motor actuators.

However, since Lashley’s seminal proposal^12^ there has been an alternative account suggesting that all elements of a planned sequence are active simultaneously before execution leading to the characteristic finding of transposition errors amongst nearby elements^13^, e.g. as observed in speech or typing. So-called ‘competitive queuing’ models can formally explain this behavior by introducing a parallel preparation layer that determines serial order by competitive interactions between sequence elements driven by differing levels of excitation in accordance with the sequence (Fig. 1a). The most active node wins the competition, generates the corresponding action and is then self-inhibited through the planning layer, allowing the next most strongly activated node to generate the next action, effectively allowing the conversion of a parallel planning code into a temporally structured serial output during execution. Crucially, it has been suggested that the respective excitation gradient may be determined through the association of each item to an independent timing or position signal that is reset prior to sequence production, such as a decaying start signal^14^, a combination of start and end signals^15^, or dynamic states for each serial position^16^. This is in contrast to RNN network models of control where serial order conjunctively codes for both position and content^17^.

Direct neurophysiological evidence in support of the competitive queuing model has been obtained by Averbeck and colleagues^18^, who taught non-human primates to copy geometrical shapes presented on the computer screen while they recorded multi-unit activity from the prefrontal cortex, an area homologous to inferior frontal cortex in the human brain^19^. Specifically, they characterized multi-unit patterns during the production of each segment in the shape-drawing sequence, and then classified the likelihood of that segment pattern becoming active during movement preparation and production. Their findings revealed that the likelihood of segment activation towards the end of the preparation phase is determined by its respective position in the sequence (Fig. 1b), as predicted by competitive queuing models. However, evidence in support of the competitive queuing model has yet to be reported in humans. Moreover, it remains unclear whether this planning process reflects a graded preparation of specific motor elements in the sequence, or instead a higher-level timing signal, which transfers across motor sequences^15,16,20,21^.

Here, we provide direct evidence for competitive queuing in the human brain during the preparation of accurately timed finger sequences from memory using non-invasive whole head recordings (magneto-encephalography; MEG), replicating previous findings from invasive electrophysiology in non-human primates^18^. Further, we demonstrate that the observed competitive queuing pattern in the MEG signal during preparation reflects effector-independent temporal sequence (serial position) planning and cannot be explained by a graded muscular pre-activation. Finally, we show that the fidelity of competitive queuing correlates with behavioural performance across participants, suggesting that this neural planning strategy is relevant for skilled sequence production and timing.

**Figure 1.**
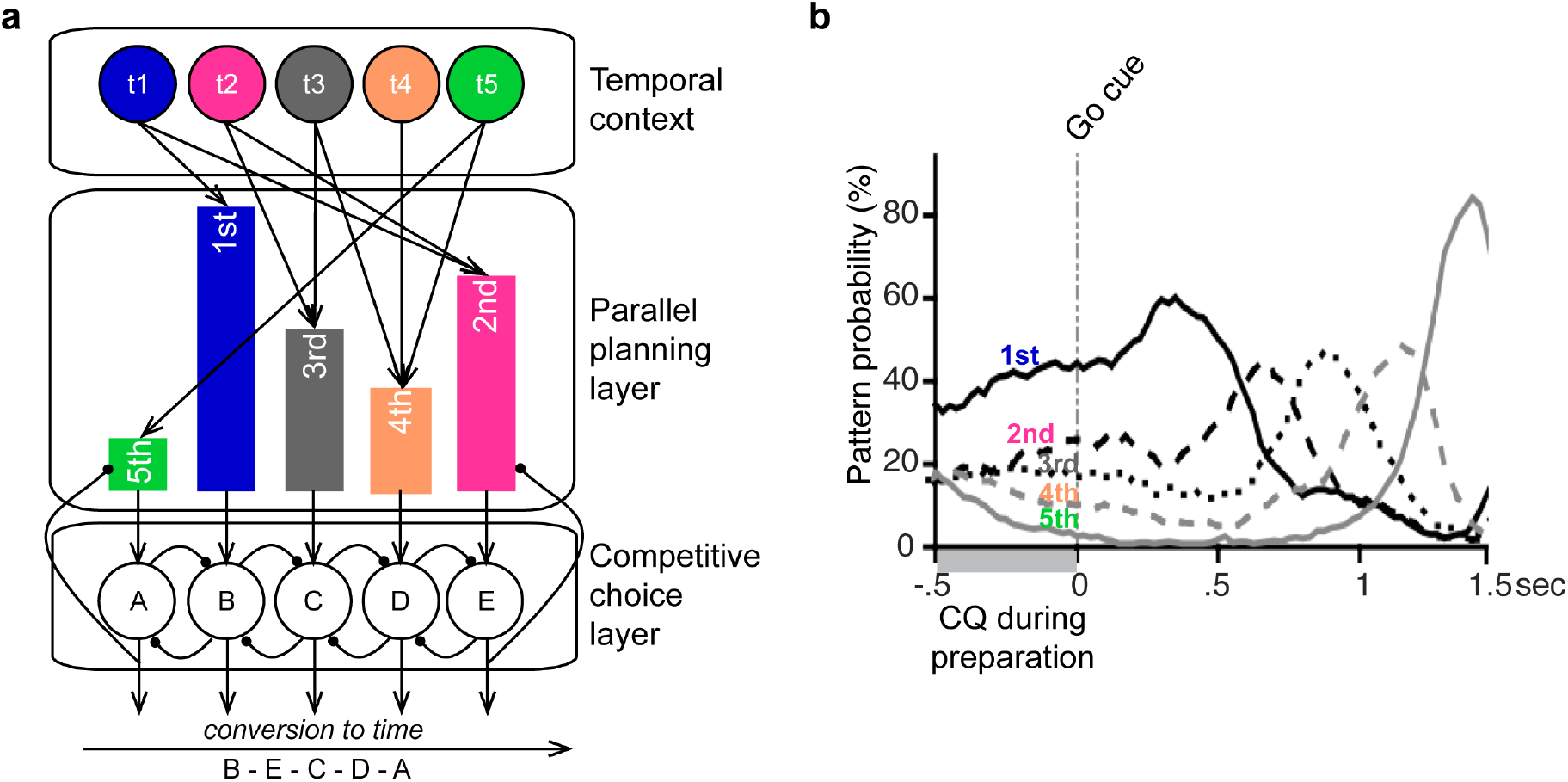
Competitive queuing model and prior data. **a.** Competitive queuing (CQ) model. The competitive choice layer contains nodes representing possible sequence items –such as muscular synergies A, B, C, D and E. Planning the temporal organization of actions to produce a sequence is implemented by activating the employed sequence items in parallel, with the highest activation allocated to the first and the lowest activation to the last sequence item, respectively. Crucially, this weight is determined by associating each item with a different state of a postulated neural “temporal context”^16^. Links from temporal context units to the item-specific parallel planning units are established using Hebbian learning. The most active node in the competitive choice layer wins the competition, generating the corresponding action and is then self-inhibited through the planning layer, allowing the next most strongly activated node to generate the next action. This iterative process allows the conversion of a parallel planning code into a temporally structured serial output. **b.** Averbeck et al. recorded the activity of individual neurons simultaneously in the prefrontal cortex while monkeys drew geometrical shapes, i.e. produced stereotyped sequences of trained movements. Each of the sequence segments was associated with distinct patterns of neuronal ensemble activity. The authors showed that 1) these patterns were present during sequence preparation and 2) the likelihood of any pattern being present during this time predicted the serial position of that movement in the motor sequence. These results are consistent with the graded parallel preparation of sequence items predicted by the competitive queuing model.

## Results

### Sequence performance shows independent temporal transfer

Participants were trained for two days to associate four abstract visual cues with the production of four finger sequences from memory following a ‘Go’ cue (Fig. 2a). Each sequence consisted of five-finger presses (finger order) and five target intervals between finger presses (timing or temporal interval order), respectively. Sequence construction followed a factorial design, with two temporal *(T1 and T2)* and two finger order sequences *(F1 and F2)* resulting in four unique temporally structured finger sequences (Fig. 2a). While in the majority of participants the production of the correct sequences was temporally aligned according to *T1* and *T2* across finger orders *F1* and *F2*, respectively (Fig. 2b), there was also substantial variation of timing accuracy between participants. Mean temporal error from target interval structure across the whole group ranged from 10-33% deviation from the target interval sequence (mean: 19%, SD: 7%; Fig. 2b). The participants were grouped using a median split of interval deviation (temporal error) for later analysis with more accurate participants showing on average 14% (SD: 3%) and the less accurate 25% (SD: 5%) absolute deviation from target interval. On average, participants tended to produce finger press intervals 7% shorter than the respective target intervals, resulting in sequences that were slightly faster than the target sequence. Subjects who were less accurate (larger absolute deviation from target) were also more likely to produce shorter, rather than longer intervals *(r* = -.817, *p* < .001, two-tailed).

To evaluate sequence-specific learning, we conducted a post-training test on the day before the MEG session during which participants were ask to synchronize their respective presses to a visual finger cue, as in the first stages of training (cf. Methods). Participants showed significantly more accurate synchronization to visual sequences when they encountered trained sequences, as well as sequences with trained finger order or trained temporal interval order to untrained control sequences (Fig. 3a-b). This result confirmed an independent representation of the temporal structure of sequences which can be utilized across finger sequences, in line with previous studies ^22-24^. Subject with more pronounced sequence-specific learning produced larger inter-press-intervals during the MEG session (r = .501, p < .001, two-tailed), suggesting that the tendency to compress the sequence was a marker of poor skill learning.

**Figure 2.**
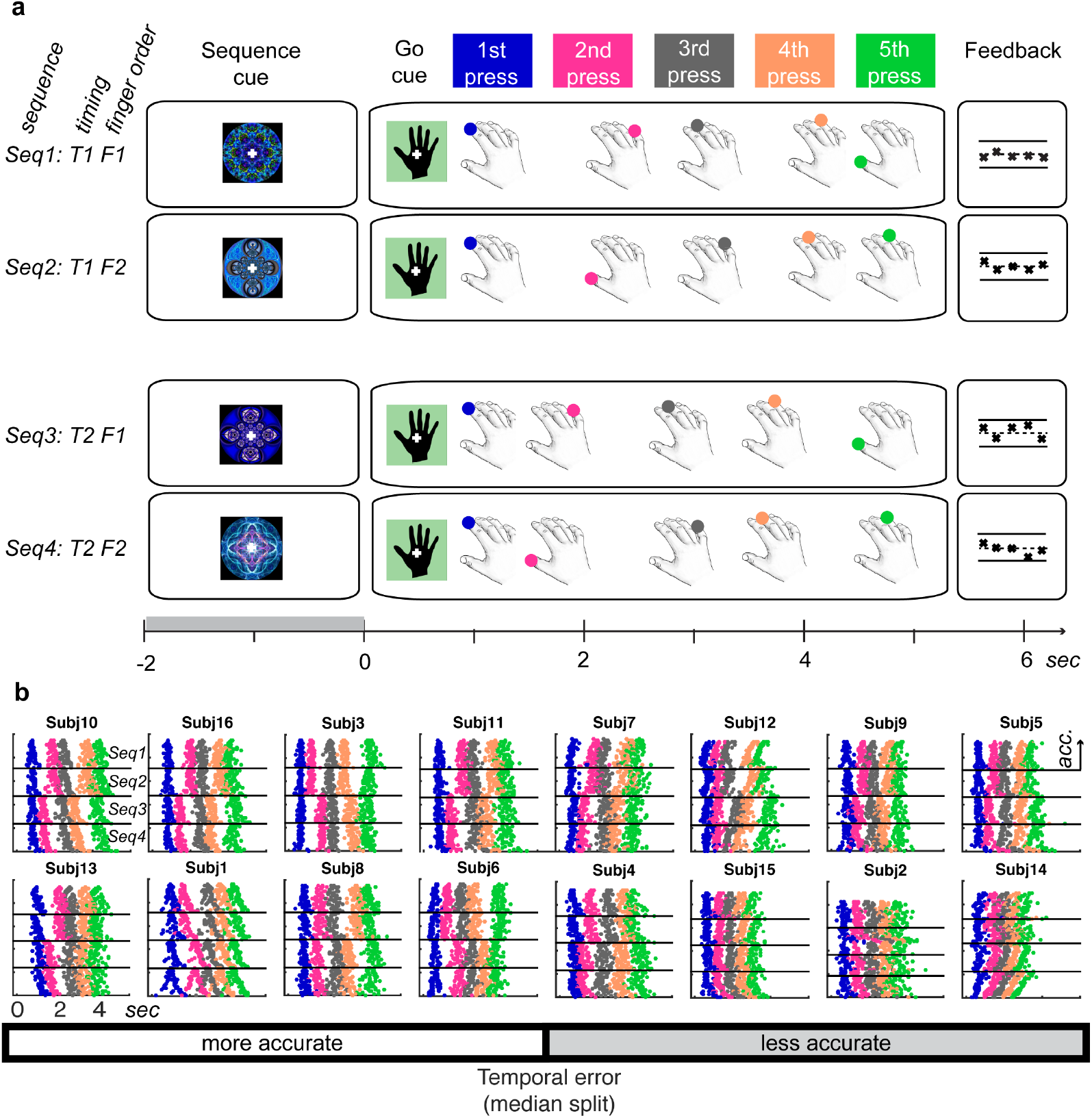
Task and production of sequences from memory. **a.** Trial design in the MEG experiment: Subjects were required to prepare and produce one of four finger press sequences as indicated by a unique visual fractal cue, followed by a ‘Go’ cue. The four sequences were unique combinations of two finger orders (F1 and F2) and two temporal or interval orders (T1 and T2) and were generated randomly for each subject. The association between randomly assigned abstract visual cues and each sequence was learned during two days of training preceding the MEG session. Each sequence started with the same target finger and target interval between ‘Go’ cue and first finger press within subjects. The ‘Go’ cue remained on the screen throughout the production period. To prevent drifts in performance and maintain attention to the task, subjects received feedback on the finger and temporal accuracy of their five finger presses after each trial with crosses (‘x ‘) signifying correct and dashes (‘-’) incorrect finger press responses across positions. The relative position of crosses on the y-axis relative to a dashed midline indicated temporal accuracy (too early being below or too late above the midline; outer lines −100 and 100 percent of target interval). b. Individual subjects ‘ raster plots show the timing of single button presses for each trial produced from memory after the ‘Go ‘ cue (t=0) during the MEG session. The colour code corresponds to the press position in Fig. 1a. The trials for each sequence are grouped by sequence number and within these groupings sorted by temporal accuracy relative to the target interval structure with the most accurate in the top to least accurate trials in the bottom rows. Subjects are grouped into more accurate and less accurate based in a median split of average temporal performance error. Here only trials with correct finger order were used for analysis. Within these subgroups subjects are arranged according to their mean temporal performance with the most accurate at the top left to the least accurate in the bottom right, respectively.

**Figure 3:**
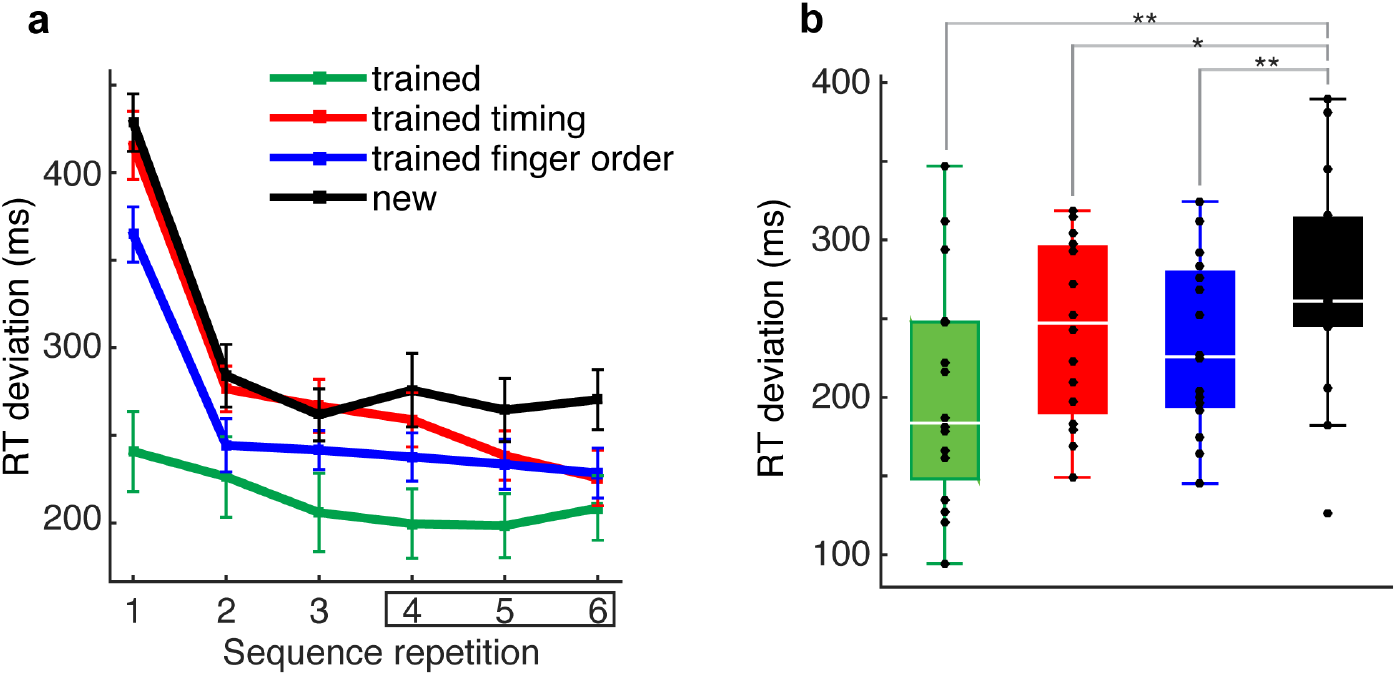
Independent transfer of timing to untrained finger sequences. **a.** A visually cued synchronization task, during which subjects had to synchronize finger presses with finger sequences presented on the screen, was used to assess sequence-specific advantages due to training. Here absolute RT deviation from finger cue (absolute RT) was determined for each condition. Repeating sequences six times in the test phase yielded an immediate decrease of absolute RT deviation from target for trained sequences (green) relative to untrained sequences (black). Sequences with a trained finger order, but an untrained timing (blue) also produced immediate benefits. Sequences with a trained timing, but an untrained finger order (red) also showed behavioural advantages, but only after the first three sequence repetitions. This is a replication of previously reported findings with evidence for a dedicated encoding of temporal structure independently of specific finger presses, as revealed by faster behavioural acquisition of new sequences compared to a control condition ^22,23^. **b.** Mean RT deviation across subjects after the first three sequence repetitions showed behavioural advantages for the trained sequence, as well as the trained finger order and trained temporal transfer conditions, compared to untrained sequences, suggesting that both the finger order and their relative timing were represented independently.** p < 0.01, *p < 0.05, one-sided t-test.

### MEG evidence for competitive queuing

During the MEG scan, finger sequences were visually cued with an abstract shape for 1. 8 - 2.2 seconds before the ‘Go’ cue. This provided participants with a short period to retrieve and prepare the corresponding sequence from memory prior to production. We leveraged multivariate linear discriminant analysis (LDA) to characterize whole head MEG activity patterns associated with the execution of each finger press. Specifically, we trained our classifier on the average MEG pattern across all sensors in a 10ms window immediately preceding each finger press during sequence production, and then applied the classifier to successive 10ms time windows during sequence preparation, in order to establish the osterior probability of each press-related pattern appearing during that period (Fig. 4).

This analysis demonstrated that, during sequence preparation, the probability of each finger press being decoded across time windows was scaled according to its serial position in the sequence (Fig. 5a-c; Supplementary Fig. 1a for individual data), analogous to the findings 18 obtained using invasive electrophysiology recordings in macaques . Specifically, the mean pattern probability during the final 1s of the preparation period was modulated by press position (F(1.71, 25.60) = 42.23, *p* < .001, *n*^2^ = 0.738; one-way repeated measures ANOVA, Greenhouse-Geisser corrected, *χ^2^* (9) = 37.56, *p* < .001). As predicted by the competitive queuing hypothesis, pattern probabilities during this period were significantly higher for 1^st^ vs. 2^nd^ (*t*(15) = 4.62, *p* < .001), 2^nd^ vs. 3^rd^ (*t*(15) = 4.57, *p* < .001), and 3^rd^ vs., 4^th^ (*t*(15) = 4.51, *p* < .001) finger presses, whilst the probability of decoding the 4^th^ press was not significantly higher than that of the 5^th^ (t(15) = 1.78, *p* = .19; one-tailed t-tests according to the competitive queuing hypothesis, Bonferroni-corrected for four comparisons).

We then examined whether competitive queuing during sequence preparation was preserved when classifying MEG patterns across sequences with the same target timing, but a different finger order. If competitive queuing reflected the upcoming order of specific effectors (here: fingers), we would expect the accurate queuing of pattern probabilities to collapse since the upcoming finger order is rearranged relative to the training order in the 2^nd^ to 5^th^ positions, i.e. the training pattern for a particular press position would reflect a different finger from that in the test pattern. In contrast, if competitive queuing during preparation was driven by the representation of ordinal position within the sequence regardless of effectors (neural patterns related to the state of being in 1^st^ to 5^th^ position in the sequence), then competitive queuing during the preparation phase should be upheld across sequences with a different finger order.

This analysis revealed that competitive queuing during the preparation period was qualitatively preserved across sequences with the same temporal interval, but differing finger order (Fig. 5a-c; Supplementary Fig. 1b and Supplementary Fig. 2 for individual data). Specifically, the pattern probability was still modulated by press position (F(1.68, 25.15) = 26.97, *p* < .001, *n^2^* = 0.643; one-way repeated measures ANOVA, Greenhouse-Geisser corrected, *χ^2^* (9) = 39.87, p < .001). Pattern probabilities at the end of the preparation period were significantly higher for 1^st^ vs. 2^nd^ (t(15) = 4.06*,p* = .002), 2^nd^ vs. 3^rd^ (t(15) = 3.29*,p* < .001), 3^rd^ vs., 4^th^ (t(15) = 3.97, *p* = .002) finger presses, whilst the probability of decoding the 4^th^ press was – once again – not significantly higher than that of the 5^th^ (t(15) = 0.98, *p* = .68; one-tailed t-tests according to the competitive queuing hypothesis, Bonferroni-corrected for four comparisons).

To determine the strength and variability of competitive queuing across participants based on the classification within sequences with the same finger order and across sequences with a different finger order, respectively, we calculated the average distance between consecutive press probabilities for each participant. Consistent with the results described above, this distance parameter was significantly above zero across participants regardless of whether the training data came from sequences with the same (‘within’) or different (‘across’) finger orders (Fig. 5c). Crucially, the average distance between press probabilities remained significantly greater than zero, even when the probability for the 1^st^ finger press (same across sequences in each participant; Fig. 2a) was excluded (2^nd^ to 5^th^ press, Fig. 5c). Moreover, across subjects, the average competitive queuing distance values obtained from the classification ‘within’ the same finger sequences were also highly correlated with those obtained ‘across’ finger sequences, regardless of whether the pattern for the first press in the sequence was taken into account (mean probability distance 1^st^ – 5^th^ press patterns: *r* = .991, *p* <.001; mean probability distance 2^nd^ to 5^th^ press patterns: *r* = .982, *p* <.001). These results confirm the manifestation of a finger-independent, and therefore, transferable positional code during the preparation of motor sequences.

In addition to the preparation phase, we also examined the dynamics of finger press probabilities during the production phase, in order to quantify subsequent modulations between ‘within’ and ‘across’ finger sequence classifications. Although the phasic execution-related peaks were markedly attenuated in the ‘across’, compared to ‘within’ sequence classification in the positions where the finger order diverged (2^nd^ - 5^th^ press; t(15) = 12.51, *p* < .001, paired one-tailed t-test), the respective press probabilities still increased above chance (20%) in the ‘across’ finger classification at the time of the respective finger presses (t(15) = 8.50, *p* < .001, paired one-tailed t-test). This suggests that the finger-independent, positional code established during the preparation period was utilized during sequence production.

However, while competitive queuing during preparation was qualitatively preserved between the ‘within’ and ‘across’ finger sequence analyses, the average distance between adjacent probabilities nevertheless decreased significantly in the latter (t(15) = 5.68, *p* < .001, paired one-tailed t-test). This advocates the loss of a conjunctive representation of ordinal structure with the respective finger during both the preparation and production phases. In sum, the above results indicate that competitive queuing during sequence preparation includes the planning of an action-independent temporal template for action sequences, in addition to the queuing of specific effectors^19^.

**Figure 4:**
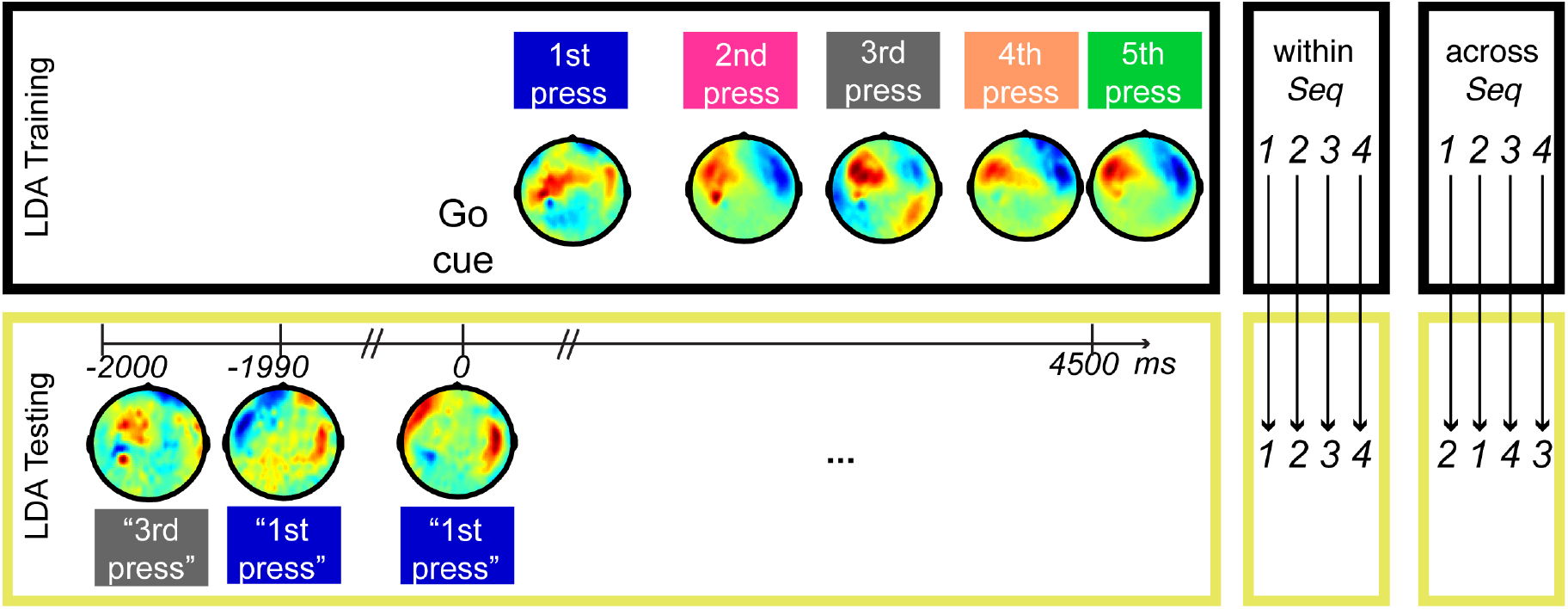
Linear discriminant analysis and classification. A Gaussian-linear classifier was trained to discriminate patterns of mean signal amplitude in a 10ms time window preceding each of the five button presses across all sensors in a data set consisting of correct trials only. Example mean signal amplitude values across the scalp are shown for a representative participant (temporal accuracy z=0.45). The test data set consisted of non-overlapping 10ms time windows starting 2000ms before and 4500ms after the Go cue. For each time window in each trial, we calculated the probability of each button press (i.e. how closely the observed activity pattern resembled that relating to each button press in the training data). For illustration purposes, only the pattern with the highest probability is shown (inverted commas) for one example trial in the same participant. To test whether the neural competitive queuing of sequence elements during sequence preparation reflected the weighting of patterns related to the ordinal structure of the sequence, rather than effector specific activity, we also trained and tested the classifier using sequences with the same timing, but a different finger order (‘across’, c*f* Fig. 2), respectively. In addition, the same analysis was performed on the mean signal amplitude from the four concurrently recorded EMG channels (not shown).

**Figure 5:**
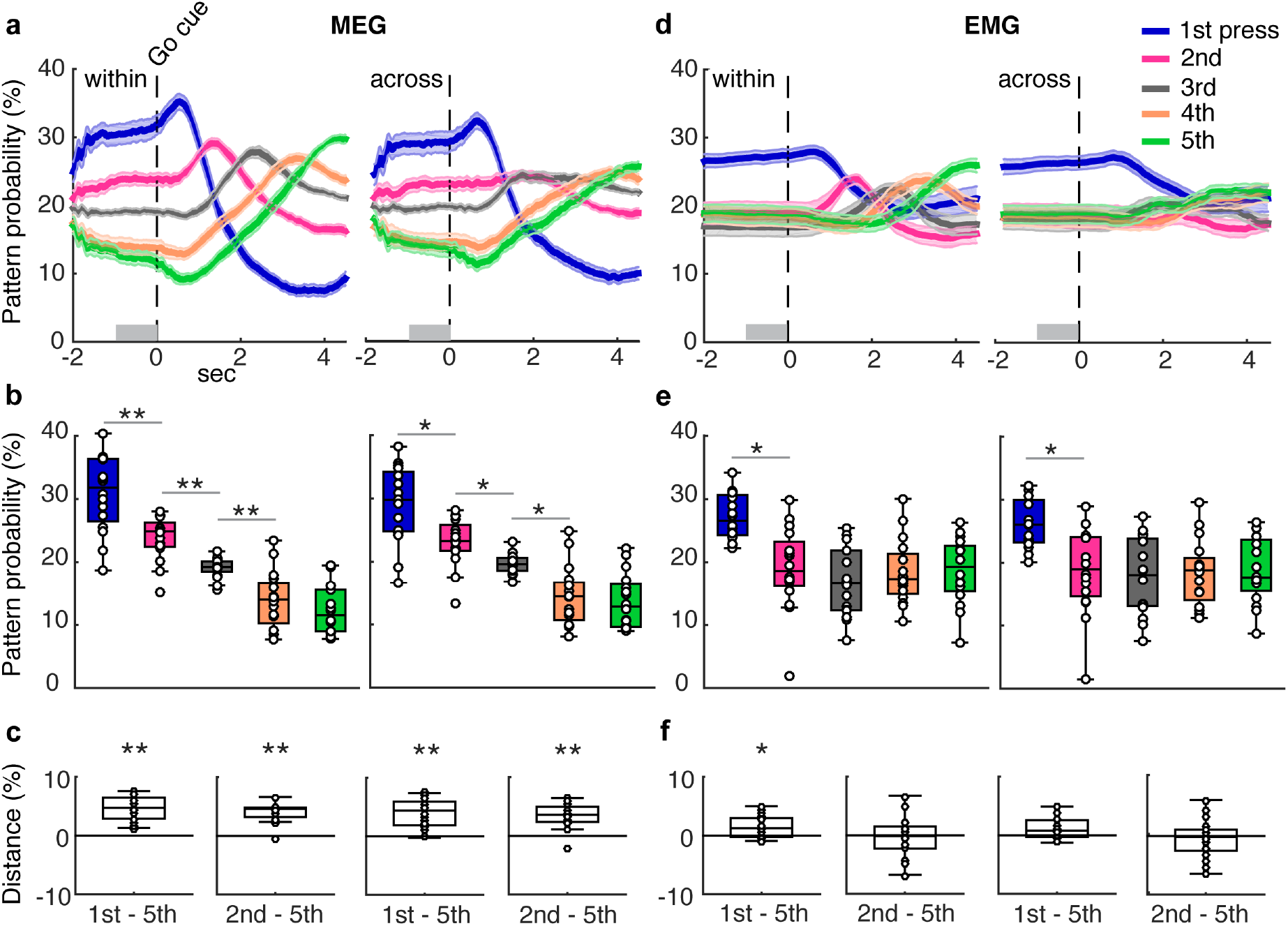
Neural vs. muscular markers of competitive queuing. **a.** MEG patterns: Trace plots display the probability of each pattern classification related to the 1^st^-5^th^ finger press 2000ms before and ending 4500ms after the ‘Go’ cue. Pattern probabilities ‘within’ refers to training and testing within the same sequence trials. Probabilities ‘across ‘ patterns refers to testing and training across sequences with the same target timing, but performed with a different finger sequence (Fig. 3). b. Boxplots show the mean probability for each press pattern and c. the mean distance between consecutive press pattern probabilities in the final 1s before the ‘Go ‘ cue. The latter include the average distance between consecutive press patterns of all (left panel) and 2^nd^-5^th^ press patterns (right panel), respectively. Note that for each subject the sequences always started with the same finger and target interval. Thus the classification ‘across ‘ different fingers applies to 2^nd^ to 5^th^ position only, providing a dissociation between finger and position- or timing-related pattern probabilities. Together the MEG results suggest a competitive queuing of sequential movement patterns which generalizes across finger sequences with the same temporal structure. d-f. EMG patterns: Results illustrating the probability of muscular patterns are displayed in the same format as panels a-c. In contrast to the MEG classification results, only the pattern for the first element in the sequence was elevated in muscular space suggesting that, at the periphery, only the first element in the sequence was prepared before the ‘Go’ cue. It also implies that the complete competitive queuing pattern in the MEG results is not driven by a corresponding graded activation of finger presses or weighted sensory feedback from the periphery. CQ: competitive queuing; ** p < 0.001, * p < .05 (t-tests against zero, Bonferroni corrected for four tests for MEG and EMG separately).

### Competitive queuing is not driven by muscle activity

Whilst the above neural signature of competitive queuing suggests a top-down signal for the temporal planning of finger sequences, it is still possible that this pattern is partly driven by a weighted activation of muscles at the periphery, such that the muscle synergy activations related to each finger movement are weighted before production according to their occurrence. Hence, using the same LDA procedure as for the MEG data training and classification (Fig. 3), we examined data obtained from muscles of the right hand (flexor carpi radialis, abductor polices brevis, abductor digiti minimi, first dorsal interossei) concurrently to the MEG recording. This analysis revealed that, during preparation, only the pattern probability for the first finger sequence press in the sequence – performed by the same finger across all four sequences – was elevated above those for the other sequence elements.

Specifically, the EMG pattern probability at the end of the preparation period was modulated by press position for both ‘within’ (F(4, 60) = 7.44, *p* < .001, *n^2^* = 0.332; one-way repeated measures ANOVA) and ‘across’ finger sequence classification (F(2.74,41.17) = 5.27, *p* = .005, *n^2^* = 0.260; one-way repeated measures ANOVA). However, in contrast to the MEG data, this effect was driven purely by the elevation of the probability of the first press in the sequence (1^st^ vs. 2^nd^ ‘within’: *t* = 3.882, *p* = .003 and 1^st^ vs. 2^nd^ ‘across’: *t* = 3.583, *p* = .005), with all other differences between adjacent press probabilities being not significant *(p* > .775, one-tailed t-tests according to the competitive queuing hypothesis, Bonferroni-corrected for four comparisons; Fig. 5e). Furthermore, the EMG distance between consecutive press probabilities during preparation was significantly above zero when the probability for the 1^st^ finger press was included in the distance calculation, but not when the only 2^nd^ to 5^th^ presses were taken into account (Fig. 5f).

Finally, the distance between consecutive press probabilities in the MEG data did not correlate with the distance in the EMG data, neither in the within (r = .275, *p* = .303), nor in the across sequence analysis (r = .286, *p* = .283). Whilst the data provides a strong indication for the muscular preparation of the first press before the ‘Go’ cue, we could find no evidence for weighted muscular synergies driving the competitive queuing pattern in the central nervous system during sequence preparation.

### Competitive queuing predicts behavioural accuracy

Finally, we asked whether the degree of neural competitive queuing during sequence preparation was relevant for the subsequent production of the sequences from memory. Specifically, we examined whether the average distance between pattern probabilities at the end of the preparation period (1s window immediately preceding the ‘Go’ cue) predicted the sum of points (reflecting finger order and timing accuracy) gained across trials in the MEG session (cf. Methods) and, more specifically, the temporal accuracy during the production period.

Our findings suggest that participants with a larger mean distance between adjacent press probabilities according to their sequence positions, tended to gain more points during the MEG session (r = .517, *p* = .020) and produce smaller temporal errors relative to the target timing structure of sequences (r = -.571, *p* = .010; Fig. 6a and b). In particular, the probability dynamics shown in Fig. 6c illustrate the striking differences in the fidelity of competitive queuing during sequence preparation between participants with more and less accurate behavioural performance, as well as the relative preservation of phasic response curves during the serial execution period. Crucially, this correlation with behavioural accuracy was also unique to MEG patterns. Despite the elevation of the first EMG press pattern probability prior to the ‘Go’ cue, which could have played a role in subsequent sequence production, the fidelity of competitive queuing in EMG patterns did not show any relationship with overall points gained or the size of temporal errors (Fig. 6e-g).

In contrast to the correlation between temporal accuracy and mean pattern probability distance across participants, we did not find evidence for the pattern probability distance during preparation to predict temporal accuracy during the subsequent production on a trial-by-trial basis. Within participants, the trial-by-trial correlation coefficients ranged from *r* = - .251 to *r* = .140 (SD = .093) with a predicted negative correlation being significant at *p* < 0.05 in only 5 out of 16 participants (Fig. 6d, grouped by timing accuracy). Accordingly, participants with a more pronounced neural competitive queuing pattern during sequence preparation had better overall performance in the sequence production phase. However, this neural signal did not guarantee high execution accuracy at the trial-by-trial level, which may be influenced by other downstream processes for motor implementation.

For the EMG patterns, in line with the group analysis, we did not find any consistent significant trial-by-trial correlations between the EMG pattern probability distance and temporal accuracy. Within participants the trial-by-trial correlation coefficients ranged from *r* = -.208 to *r* = .294 (SD = .137) with a predicted negative correlation being significant at*p* 0.05 in only 3 out of 16 participants (Fig. 6h, grouped by timing accuracy). Therefore, in the case of EMG data, there is no evidence of the elevated probability for the pattern related to the first press to be associated with performance, neither across nor within participants (trial-by-trial).

**Figure 6:**
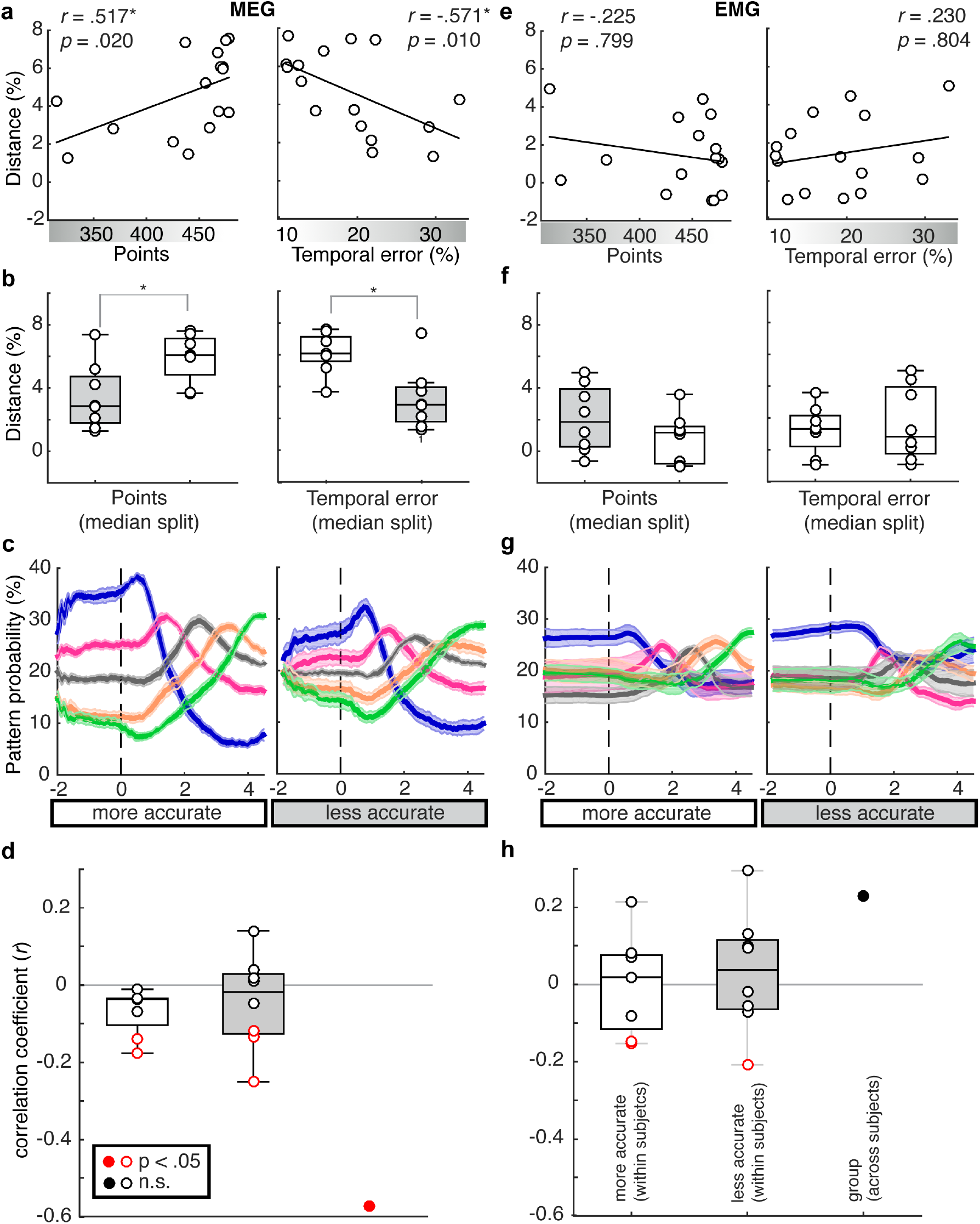
Correlation with behavior. **a.** Overall points acquired during the MEG session correlated positively (left panel) and the temporal error from target intervals (right panel) correlated negatively with the average distance between consecutive MEG probability patterns extracted from the end of the preparation period in the ‘within ‘ sequence classification (1^st^ – 5^th^) as a proxy of the strength of competitive queuing. b. The median split based on points (finger order and timing accuracy) and temporal error revealed a significantly stronger competitive queuing for subjects with more points and lower temporal error (higher temporal accuracy). Again this effect could not be observed for the EMG patterns, despite the fact that the elevation of the probability for the first element in the sequence was enhanced compared to the other patterns and might have contributed to performance. c. Trace plots as in Fig. 4a grouped according to the median split by temporal error. Subjects with a more precise temporal sequencing showed a higher spread of press related pattern probabilities, suggesting that more pronounced competitive queuing is associated with superior performance. d. The correlation of temporal error and distance between press patterns in the preparation period was driven by differences between subjects, rather than by trial-to-trial fluctuations. Trial-by-trial pattern distance probabilities and temporal performance were correlated in four out 16 subjects. This finding suggests that the strength of neural competitive queuing during sequence planning predicted overall, but not trial-to-trial performance in the subsequent execution period. e-h. Association between pattern probability spread and behavior did not hold up for the EMG patterns.4 ^**^ p < .01, * p < .05, one-sided t-test and linear correlations in line with the directional hypothesis for behavior and competitive queuing patterns. Two-sided t-tests and correlations were also significant at p <.05.

### Competitive queuing originates from parahippocampal and cerebellar sources

Next, we sought to identify the neural origins of competitive queuing during preparation in MEG sensor and source space. To this end, we made use of searchlight analysis to determine where the competitive queuing signal observed during the preparation period was most pronounced. Specifically, we quantified the average distance between consecutive press probabilities at a single time point 0.5s before the ‘Go’ cue in each trial, after training and testing our classifiers on data from the same finger press sequences (‘within’ sequence analysis). In addition, these average distance values were z-transformed across all sensor and voxel searchlights within each participant prior to group level statistical analysis. The most pronounced effect of competitive queuing during preparation appeared in the right temporal sensors (*t*(15) = 3.58, *p* < 0.01, uncorrected, Fig. 7a). In line with this finding, the same analysis conducted in source space showed a significant competitive queuing distance effect to originate from a single cluster (*p_c_l_uster_* < 0.001) comprising the right temporal cortex, specifically the parahippocampus, extending ventrally into the fusiform area (MNI coordinates of peak voxel: 30, −30, −24, *t(15)* = 6.94, *p* = .005, p-value FWE-corrected at voxel level); and the right (ipsilateral) cerebellum, specifically lobules VIII (MNI coordinates of peak voxel: 34, −30, −50, *t(15)* = 6.51, *p* = .009) and V (MNI coordinates of peak voxel: 64, −46, −30, *t(15)* = 5.72, *p* = .046, Fig. 7c).

In addition, to localize the sources of the press-related training patterns - that is of conjunctive representations of position and effector observed during sequence production – we trained and tested our classifier on MEG data obtained immediately prior to each finger press within each motor sequence using a five-fold cross-validation procedure. As before, decoding accuracy values were z-transformed across sensors and voxel searchlights within each participant prior to group level statistical analysis. Consistent with prior findings^25,26^, the accuracy of finger press decoding was most pronounced in the sensors located above the left sensory and motor areas contralateral to the moving hand and the right temporal sensors (t(15) = 3.11, p<0.01, uncorrected, Fig. 7b), the latter partly containing the same significant sensors as in the competitive queuing analysis from the preparation period. At the source level, we found a large significant cluster (*p_cluster_* < 0.001) comprising contralateral primary sensory and motor regions with peaks in the primary sensory cortex (MNI coordinates of peak voxels: −22, −38, 52, t(15)=11.19, p<0.001 and −54, −30, 40, *t*(15)=8.87, p=0.001), extending medially into the cingulate cortex (MNI coordinates of peak voxel: −14, −28, 44,ŕŕ75j=8_ğ_96,/?=0_ğ_001; Fig. 7d)_ğ_

Finally, to gain a more detailed picture of the complementary contributions of these regions during sequence preparation and production (Fig. 7a-d, line and box plots), we examined competitive queuing during the preparation phase and finger press decoding accuracy from the production phase in each group of significant sensors and peak voxels of significant clusters reported above. Interestingly, finger press decoding accuracy in the right temporal sensors, as well as the right parahippocampal and cerebellar sources identified by the competitive queuing analysis during the production phase, showed above chance finger press decoding accuracy in all sixteen subjects (p<0.001, one-sided t-tests against chance level). This suggests that activity patterns from these sources also contained information about the finger press being executed during sequence production, although the size of the effect was eclipsed by the concurrent representation of finger press identity in contralateral S1/M1 regions. In contrast, the probabilities in central sensors, as well as contralateral S1/M1 during preparation did not show any evidence for a competitive queuing gradient (sensor level: one-way repeated measures ANOVA, F(2.32, 17.77) = 0.37, *p* = .72, *η^2^* = 0.024; Greenhouse-Geisser corrected, *X* (9) = 21.58, *p* = .01; source level: F(3, 32.18) = 1.82, *p* = .16, *η2* = 0.108; Greenhouse-Geisser corrected, *X* (9) = 18.71, *p* = .03), suggesting that contralateral neocortical sensorimotor areas did not contribute to establishing the temporal order of finger presses before execution. In sum, our results indicate the special role of the parahippocampal and effector-related cerebellar sources in establishing a competitive queuing gradient during sequence preparation and its utilization during sequence execution, whilst contralateral primary sensorimotor sources appear to contribute to the task during sequence execution only.

**Figure 7:**
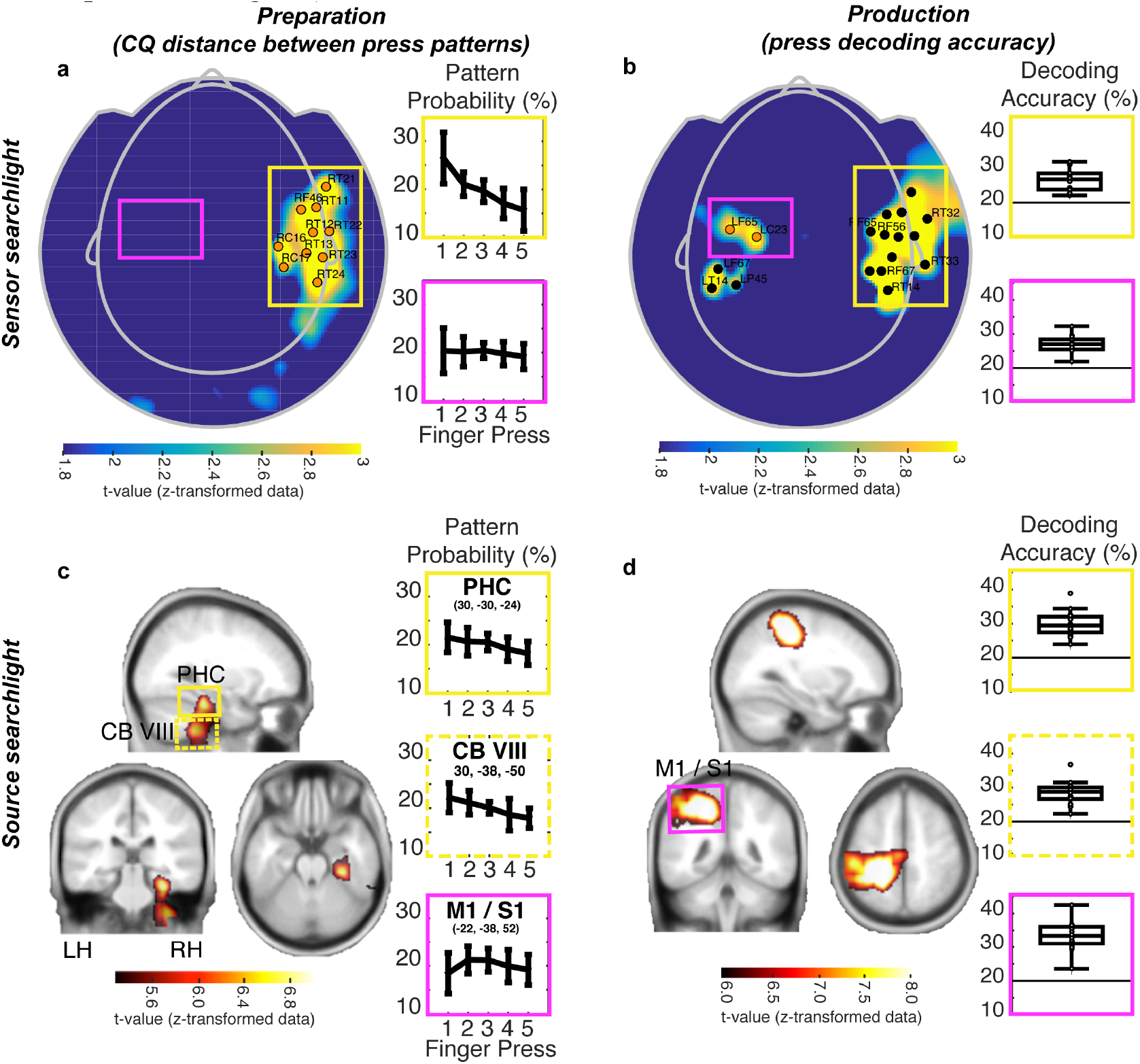
Significant sensors and sources with most pronounced competitive queuing (CQ) strength during preparation. **a.** CQ distance during preparation (sensor level): A sensor level searchlight analysis demonstrated that competitive queuing during preparation (0.5s before the ‘Go’ cue) was most pronounced in the right temporal sensors (p<0.01, uncorrected). Line graphs show corresponding pattern probabilities during preparation from significant sensors of interest marked in orange in panels a and b and associated with right temporal (yellow) and left sensorimotor sites (magenta). Sensorimotor sites showed no evidence of competitive queuing during sequence preparation. **b**. Decoding accuracy offinger presses during production (sensor level): Searchlight analysis revealed that sensors driving the differences between the press patterns during production were located above the left (contralateral) sensorimotor sensors, in addition to right temporal sensors. Boxplots show individual decoding accuracies in sensors of interest and suggest that temporal sensors remained relevant during sequence production. **c**. CQ distance during preparation (source level): The combined CQ distance searchlight analysis at the source level (corresponding to sensor level data in panel a) revealed the right parahippocampal area and right (ipsilateral) cerebellar lobules V and VIII as likely sources of competitive queuing during preparation. Data is plotted as a t-statistic at a threshold of t(15) > 5.24, p < .05 (FWE corrected at the voxel level) centered at the peak voxel (MNI coordinates: 30, −30, −24). Line graphs show corresponding pattern probabilities during preparation from peak voxels in significant clusters identified in the respective analyses presented in panels c and d, specifically right parahippocampal (yellow), right cerebellar lobule VIII (yellow dashed) and left sensorimotor neocortex (magenta). As in panel a, sensorimotor sites showed no evidence of competitive queuing during sequence preparation. **d**. Decoding accuracy of finger presses during production (source level): Searchlight analysis revealed that the differences between the press patterns during production were driven by the left (contralateral) sensorimotor neocortex. Data is plotted as a t-statistic at a threshold of t(15) > 5.93, p < .05 (FWE corrected at the voxel level) centered at the peak voxel (MNI coordinates: −22, −36, 52). Boxplots show individual decoding accuracies in peak voxels of interest as in panel c, suggesting that despite not being significant at the whole brain level, temporal sensors remained relevant during sequence production, with above chance (20%) decoding accuracies. V: lobule V; VIII: lobule VIII; CB: cerebellum; CQ: competitive queuing; LF: left frontal; LH: left hemisphere; LP: left parietal; LT: left temporal; M1: primary motor cortex; PHC: parahippocampal area; RC: right central; RH: right hemisphere; RT: right temporal; S1: primary sensory cortex.

## Discussion

Using non-invasive neurophysiological recordings (MEG) in combination with multivariate pattern classification (LDA), we provide direct evidence for parallel competitive queuing of planned sequential actions in humans, in line with prior invasive electrophysiology data from non-human primates^18^ and computational models^18,3,15,17,21,27^.Our study extends these findings by showing that competitive queuing provides an abstract template for serial order which is transferable across movement sequences. Crucially, we demonstrate that the overall strength of competitive queuing during planning predicts the participant’s skill level during production, implying that this neural mechanism may be beneficial for sequence control. These findings are likely to have significance for the training of skilled sequences across diverse domains including handwriting, typing, speech, musical and athletic performance, as well as for non-invasive brain-computer interfaces for skilled motor control.

### Neural competitive queuing in humans

Despite differences in methods and species, our results bear remarkable resemblance to prior electrophysiology data obtained from macaques^18^. Specifically, our data suggests that the elements which constitute a movement sequence are prepared in parallel, with the corresponding pattern probabilities being weighted by their respective position in the subsequent sequence. After the ‘Go’ cue, these pattern probabilities transitioned into phasic increases reflecting the serial execution of the finger presses. Importantly, there was no evidence to suggest that the parallel queuing of sequential elements in the MEG may be explained by a graded activation of muscle synergies related to each finger press, as the EMG pattern showed the preparation of the first finger press in the sequence only.

Our data is at odds with an alternative model of motor sequence control which has dominated the field in the last decade, implying that skilled sequences such as handwriting or tapping sequences (e.g. Morse code production) are controlled by serial state-space trajectories in recurrent neural networks (RNN)^10,11^. Tasks exhibiting markers of competitive queuing or a dedicated encoding of sequence timing independently of the specific movements, such as the production of piano-like finger movement sequences, the drawing of geometrical shapes and other forms of skilled sequence production in humans (typing, speech) and animal models (birdsong) ^22-24,28,29^, cannot be easily reconciled with the idea of a serially evolving spatio-temporal neural trajectory in such networks. An exception is the superpositional coding of sequence elements at the onset of sequence recall produced using a RNN model by Botvinick and Plaut^17^. Their model was able to produce population activity patterns relating to each position in the sequence and exhibit a summation of these patterns at the onset of the sequence without an explicit competitive queuing architecture, rendering the RNN encoding effectively “parallel” and thus fully compatible with both the probability gradient for sequence elements during preparation shown by Averbeck et al.^18^ and our findings (Fig. 5a). Notably, the serial position was inextricably linked to the item (here: finger), which diverges from the current findings suggesting an abstract positional coding in addition to a conjunctive representation of position and finger.

It is possible that both a serial chaining and parallel queuing of sequence representations co-exist in the nervous system. Accordingly, their utilization may be determined by the kinematic features of the movement sequence itself. Specifically, in contrast to the current discrete movement sequence (involving temporal gaps in between movements), continuous and overlapping movement sequences are prone to be encoded as integrated synergies, with movement timing emerging from state-dependent control, e.g. position or velocity-dependent information ^30-33^. Notably, most skilled actions in everyday life such as speech, handwriting and tool-use involve a wide range of discrete and continuous movements necessitating the combination of these two control modes. Here future studies need to establish how different sequence types employed in these everyday domains are segmented and prepared at the neural level.

### Competitive queuing reflects finger-independent temporal planning of sequences

We assessed whether the competitive queuing pattern during preparation was driven by a) the preparation of specific movements in the sequence, or b) by an action-independent temporal template for the upcoming sequence, as predicted by models of serial recall within the framework of competitive queuing^15,16,21,34^. To this end, the classifier was trained and tested on sequences with the same target timing, but a finger order which diverged from the 2^nd^ press onwards, (Fig. 4). Our data indicate that the competitive queuing signal during preparation is largely preserved under these circumstances, whilst the phasic increases in finger press probability observed during sequence production are distorted after the effector patterns diverged. This indicates that the training patterns, corresponding to mean activity in a 10ms period before each registered finger press, contained information on the temporal position in the sequence in addition to the effector information, and that this information on the timing of sequence elements was retrieved from memory in the sequence planning stage.

The retrieval of an abstract temporal template during movement planning is in line with models suggesting that the temporal evolution of the sequence is established through the association of a dedicated timing control signal to a parallel planning layer ^15 16 2135^. Furthermore, independent behavioural transfer of trained sequence timing to new finger orders in current and previous studies^23,24,36,37^ supports the notion that temporal sequence features were encoded separately from the specific movements (Fig. 3). RNN models have been shown to acquire superpositional coding of the list elements activated at the onset of recall, however, they have not been able to account for coding independently of the item^17^. RNN models are, in principle, well-suited to serve as neural clocks^38^, and therefore as a dynamic or timing control signal one step removed from the item-specific parallel planning layer. The reported competitive queuing of sequence element probabilities before movement production could reflect a gradient of superimposed patterns encoding each ordinal position or a temporal structure in the sequence. Effectively, this mechanism implies the reinstatement of a temporal template during sequence preparation which can be utilized across different movement sequences.

We found that the most pronounced pattern of competitive queuing during preparation originated in the right parahippocampal area. Neuroimaging studies have shown the recruitment of hippocampal and parahippocampal regions during motor sequence learning tasks, specifically those involving sequences of discretely timed movements including those learned implicitly^39-41^. Moreover, parahippocampal activity has been reported to correlate specifically with the accuracy of temporal tapping patterns involving a sequence of short and long intervals^40^. Our data suggests that this parahippocampal involvement is related to the neural gradient reflecting the ordinal position of elements in a sequence with the latter facilitating accurate sequence timing. Crucially, we were able to uncover the evolution of this mechanism across preparation and production phases, demonstrating the reinstatement of the full sequential plan before sequence execution. Our results substantiate the broader involvement of hippocampal and parahippocampal areas in the timing of sequences^42,43^, and more specifically in the temporal succession or ordinal structure of sequences^44,45^.

Further, the source reconstruction indicated the involvement of ipsilateral cerebellar lobules V and VIII which have been shown to hold sensory and motor representations of fingers of the ipsilateral hand in humans ^26^. This suggests that the cerebellum works in concert with parahippocampal areas to achieve the queuing of actions during sequence preparation. Specifically, we speculate that while the parahippocampal areas may retrieve a more abstract temporal template for the sequence^44^, the representation of the actions themselves may be set by the effector-specific cerebellar circuits in the form of a fine-grained spatio-temporal forward model of the finger sequence before the onset of the production phase^46,47^. Notably, activity attributed to the parahippocampal area and the cerebellum showed above chance accuracy in decoding the sequence presses during the production period, whereas the reverse – neocortical sensorimotor regions showing competitive queuing of finger press patterns during preparation – did not hold true. This dissociation demonstrates that the structure of the upcoming sequence is pre-specified outside the neocortical regions that generate the movements and continues to be utilized during production (corresponding to temporal context and parallel planning layers in Fig. 1a), whilst the regions representing the movement synergies is involved during production only (items in the competitive choice layer in Fig. 1a).

Taken together, our data suggests that the brain learns the task structure and factorises behavior into specific actions and their general temporal struture (here sequential order). Associating actions with a sequential position and passing that position through a competitive filter may be sufficient to generate skilled sequence production. This factorisation makes learning more efficient than having to generate a unique representation of the temporal structure for each new sequence and allows for transfer of a learned structure to new motor sequences.

Our findings provide a strong case for a preparation of effector-unspecific temporal template of a sequence, however, the current study does not allow us to distinguish between the preparation of sequence position (rank-order) versus of a more fine-grained temporal planning. Future studies should manipulate the temporal structure of the sequences consistently across participants to probe how it affects the distances between the press pattern probabilities during sequence planning.

### Strength of competitive queuing during preparation predicts performance

Participants who achieved a higher skill level in the sequence production from memory showed a larger separation between adjacent press-related pattern probabilities during sequence preparation such that the competitive queuing pattern were more prominent in participants with a better overall performance (Fig. 5a-c). This is in line with the architecture of competitive queuing models, in which a more pronounced weight difference in the parallel planning layer (Fig. 1) is beneficial for sequence accuracy due to a more pronounced excitation of the proximal and inhibition of the distal elements in the sequence. Notably, despite the consistent elevation of the probabilities for the 1^st^ press during sequence planning, this effect was not paralleled by the muscular pattern data (Fig. 5d-f). Thus, it is unlikely that the MEG patterns during sequence preparation were driven by a graded muscular preparation of the sequence presses.

Remarkably, the significant correlation between overall performance and mean competitive queuing strength contrasted sharply with the absence of a trial-by-trial correlation within participants. This dissociation is a possible indication that the strength of competitive queuing during sequence preparation is a neural strategy adopted by - on average - more skilled performers. While this neural strategy may be beneficial, it does not guarantee accurate performance on each trial, possibly due to the modulation of the motor output by downstream mechanisms occurring between sequence planning and motor implementation. Although this dissociation between within and across participant correlations was striking, we cannot exclude that the absence of a trial-by-trial correlation with performance may be due to a low signal-to-noise level or that the linear classification approach (LDA) taken may not optimally capture the decision boundary between the press patterns to ensure accurate probability estimations at the level of individual trials.

The current non-invasive measure of competitive queuing during sequence planning in humans, as well as the link between overall competitive queuing strength and performance provides a promising step towards the identification of markers for skilled sequence preparation. Determining the amount of separation between sequence elements before movement production may become valuable in the context of neuro-feedback training with the aim to improve performance in patients with higher order motor impairments affecting sequence initiation and fluency, e.g. stuttering, dyspraxia and task-specific dystonia (musician’s dystonia and writer’s cramp).

Finally, our approach has the potential to advance the development of brain-machine interfaces for paralyzed patients by allowing non-invasive read-out of upcoming sequences. Previous studies have looked at predicting single-targeted movements^48^, or sequences up to two movements at a time, e.g. from invasive recordings in the premotor cortex during sequence planning^49^. The current analysis may assist the identification of a five-element sequence during the sequence planning period using EEG/MEG signals with the aim of achieving a more fluent control of external devices for skilled sequence production, such as a virtual keyboard for communication or tool use via an intelligent prosthesis.

## Conclusions

Using a finger movement task in combination with non-invasive whole-brain MEG in humans, we replicated the seminal neurophysiological findings on the parallel competitive queuing of sequence elements before sequence production in non-human primates^18^. Our data extends previous work by demonstrating that the competitive queuing of sequence presses during preparation reflects top-down planning of ordinal position or finger-independent timing and cannot be explained by a graded weighting of concurrently recorded muscular activity. More generally our data suggests that the brain learns the general structure of a motor sequence by associating actions with a transferable temporal position in the sequence, effectively factorizing the neural representation of behavioural sequences. Importantly, our findings suggest that the overall strength of competitive queuing during preparation determines the accuracy of sequence production, opening up new avenues for the rehabilitation of patients with impaired initiation and fluency of sequence production.

## Methods

### Participants

Sixteen right-handed healthy adults (9 females; mean age 24.4 years, SD: 4.9) with normal or corrected-to-normal vision participated in this experiment which included two days of training and one MEG session. Five additional participants participated in the study but had to be excluded as follows: two participants due to poor performance at the end of the training session or during the MEG session (error rate > 40%), one participant due to the absence of sequence-specific learning at the end of training, one participant due to technical issues with the MEG system and one due to discomfort during the MEG session leading to the termination of the experiment. All participants gave written informed consent to participate. The study was approved by the University College London Research Ethics Committee for Human-Based Research (UCL Ethics ID: 1338/006, Data Protection: Z6364106/2011/10/25). All participants were financially compensated for their participation.

### Experimental design

Stimuli were presented via a digital LCD projector (brightness = 1500 lumens, resolution = 1024 × 768 pixels, refresh rate = 60 Hz) onto a screen (height = 32 cm, width = 42 cm, distance from participant = 70 cm) that was parallel to the participant’s face inside a magnetically shielded room using the Cogent (www.vislab.ucl.ac.uk/cogent.php) toolbox running in MATLAB (The MathWorks, Inc., Natick, MA). Participants had a similar setup during behavioural training involving the same response device (5 buttons, Current Designs),with the visual stimuli shown on the computer screen directly and the participants being seated at an office desk. Stimuli were symmetrical abstract visual fractals. For each participant the four sequence cues were randomly selected and assigned to the sequences from a pool of sixteen fractals. Participants were presented with a feedback screen after each trial showing the number of cumulative points across the whole experiment, as well as feedback on whether they pressed the correct finger at the correct time. Figure 2 provides further details on the trial design. Participants received two points per trial for a correct finger sequence with a temporal deviation from target timing of less than 30%, one point for a correct finger sequence with a temporal deviation of less than 60% and zero points in any other case. During the first two training days participants were presented with an auditory sound concurrently with the feedback screen, which indicated 0-2 points. No auditory feedback was presented during the MEG session on day 3.

Training on day 1 consisted of 7 instructed blocks containing only trials in which the sequence was cued by a circle appearing on the target finger at the target timing after the ‘Go’ cue to which the participants had to synchronize (168 trials), 7 mixed blocks (56 instructed and 112 from memory trials following each other in a blocked 1:2 pattern, respectively) and 7 blocks with sequences produced from memory (168 trials). Training on day 2 consisted of 7 mixed blocks as previously and 12 blocks (288 trials) produced entirely from memory. Additional testing that included trained and untrained sequences (consisting of untrained finger and interval orders) presented as instructed trials was also conducted before and after training blocks on day 1 and 2, respectively. These included trained and untrained sequences presented as instructed trials. The MEG session consisted of 2 mixed blocks (16 instructed and 32 from memory trials) for participants to refresh their memory of the sequences and become accustomed to the MEG environment. This was followed by 10 blocks with concurrent MEG and EMG recordings containing trials with sequences produced entirely from memory (giving a total of 240 trials split evenly between the four trained sequences).

### Behavioural data analysis

To determine temporal accuracy in trials produced entirely from memory, we calculated a mean absolute deviation from target interval structure for each trial expressed as percentage of the target intervals. For the pre and post-test which consisted of instructed trials only, we calculated the absolute reaction time deviation from the finger cues to which the participant had to synchronize.

### MEG recordings

MEG was recorded continuously at 1200 samples/second using a whole-head 275-channel axial gradiometer system (CTF Omega, VSM MedTech) while participants sat upright in a magnetically shielded room. Head position coils were attached to nasion, left, and right auricular sites to provide anatomical co-registration.

### EMG recordings

Participants were also fitted with four EMG electrodes to measure finger movement-related muscular activity. The electrodes were placed above the flexor carpi radialis (FCR), abductor polices brevis (APB), abductor digiti minimi (ADM), first dorsal interossei (FDI). FCR was recorded with a belly-belly montage, APB, ADM and FDI with a tendon-belly montage.

### MEG and EMG preprocessing

MEG data analysis made use of SPM8^50^ (Wellcome Trust Centre for Neuroimaging, London, United Kingdom), Fieldtrip^51^ (Donders Institute for Brain Cognition and Behaviour) and custom MATLAB code. MEG data was downsampled to 1000 Hz, epoched for preprocessing into long trials spanning −2.8 to +12s around the fractal cue to include a baseline fixation, fractal cue, ‘Go’ cue, sequence production and feedback. A 48-52Hz stopband filter was then applied to remove the 50 Hz power line noise within these long epochs. Channel artefacts were inspected in each participant, but no channels were identified as corrupted in any of the datasets. Due to the need to retain as many trials as possible for pattern classification and the involvement of long epochs, physiological artifacts related to heart rate, eyeblink and breathing were identified based on the characteristic topography and time-course for each participant using ICA (RUNICA algorithm) and removed from the dataset. This procedure was carried out blind to the sequence conditions using the Fieldtrip component data browser which allows the inspection of the topography and trial-by-trial time-course of the components. ICA-corrected data was then submitted to multivariate classification analysis, with the test probabilities calculated for a shorter epoch encompassing the preparation and production phases only (−2 to 4.5 sec after the ‘Go’ cue). The EMG data was downsampled, epoched and filtered in the same way as the MEG data. No trials were removed from the data set, so that the EMG data from the same trials as in the MEG dataset was submitted to multivariate classification analysis.

### Pattern classifier analysis

First we trained a standard linear discriminant analysis (LDA) classifier to distinguish the MEG activity at the onset of each of the five button presses in a sequence. Following preprocessing, mean signal amplitude on each sensor during the 10ms period immediately before the onset of each response button press in the sequence was determined for all correct trials (mean proportion of correct trials across participants: 97.3% (SD: 3.1%); range: 87.299.7%;). These mean signal amplitude values were used as a training dataset for all correct trials of each of the four sequences, respectively. The mean sensor pattern for each button press, and the common sensor-by-sensor co-variance matrix was determined from the training data set. A Gaussian-linear multi-class classifier (cf. Supplementary Material) was then used to calculate the posterior probability of an activity pattern belonging to each of the five presses in the sequence across non-overlapping 10ms time windows in each trial of a) the same sequence (‘within’) and b) a sequence with the same target timing, but a different finger order (‘across’). The same procedure was used for the analysis of EMG patterns.

For statistical analysis of competitive queuing during sequence preparation, an average probability for each of the five press patterns was determined for per trial in a 1s time window immediately before the ‘Go’ cue. We then quantified the competitive queuing strength in each trial by taking the median difference in posterior probabilities for each pair of consecutive finger presses (i.e. from 1^st^ to 2^nd^, 2^nd^ to 3^rd^, 3^rd^ to 4^th^ and 4^th^ to 5^th^). Since the 1^st^ press had a special status with the finger identity and target onset timing remaining the same across sequences for each participant, we also calculate this distance measure without the 1^st^ press probability (median difference from 2^nd^ to 3^rd^, 3^rd^ to 4^th^ and 4^th^ to 5^th^). To display pattern probability dynamics, probabilities for each finger press pattern were averaged across trials for each 10ms time window, and then averaged across sequences. For display purposes only, the data was smoothed using a sliding average boxcar of 100 ms (i.e. ten 10ms time bins) using the MATLAB fastsmooth function (https://uk.mathworks.com/matlabcentral/fileechange/19998-fast-smoothing-function).

### Source reconstruction

The linearly constrained minimum variance (LCMV) beamformer spatial filter algorithm from the Dynamic Analysis in Source Space (DAiSS) toolbox for SPM12 was used to estimate cortical activity on a 10mm grid for whole brain analyses based on preprocessed MEG data. Co-registration to MNI coordinates was based on nasion, left and right pre-auricular fiducial points, and the forward model was derived from a single shell fit to the inner skull surface of a canonical T1 image. This data was then submitted to searchlight analysis.

### Searchlight analysis

To identify the neural origin of the competitive queuing signal during sequence preparation at the sensor and the source levels we used a searchlight approach in combination with LDA. The searchlight size corresponded to ~1% of the data – three sensors at the sensor level and 20 voxels at the source level. For the focal LDA analysis at the sensor level, we first determined the two nearest neighbours of each sensor based on the MEG sensor layout. As in the case of the whole-head analysis, the LDA classifier was then trained on mean signal amplitude from those three sensors during the 10ms period immediately before the onset of each physical button in correct trials. Next, we tested the classifier on mean signal amplitude from the same three sensors in a single 10ms time window 500ms before the ‘Go’ cue and quantified the probability of each finger press being decoded in each trial. Finally, as in the previous competitive queuing distance analysis, we computed the median distance between the probability of the 1^st^ to 5^th^ (‘within’) and 2^nd^ to 5^th^ (‘across’) finger presses, respectively, averaged across trials for each sequence and then across sequences for each participant. We then z-transformed the median distance values across sensors for each participant and computed a one-sample t-statistic on z-score values at each sensor across participants. This data was then plotted as t-statistic at the scalp level and significant sensors at p<0.01 marked with a point and a label.

The same analysis was applied to mean signal amplitude in source space using searchlights of 20 voxels as features in the LDA. Whole brain results were subjected to a random effects analysis with an uncorrected threshold of t(15) > 3.73, p<0.00. Peak voxels and t-values of significant clusters were then reported, and the data plotted as t-statistics with a threshold of t(15) > 5.24, *p* < .05 (FWE corrected at the voxel level) centred on the peak voxel ^52^. Peak voxels and t-values of significant clusters were then reported in the results and the data plotted as a t-statistic at the scalp level at a threshold of t(15) > 5.24, *p* < .05 (FWE corrected at the voxel level) centred at the peak voxel (MNI coordinates: 30, −30, −24). The data thresholded at t(15) > 3.73, *p* < .001 (uncorrected) can be found in Supplementary Figure 3 a.

Finally, to identify regions that were driving the differences between finger presses during the sequence production period, we employed a searchlight approach in combination with LDA using data from the production period only. As in the analysis described above, the searchlight size corresponded to ~1% of the data - three sensors at the sensor level and 20 voxels at the source level. Here the LDA classifier was trained to distinguish between the activity patterns of sensors or voxels during the 10ms period immediately before the onset of each physical button in correct trials. We used a five-fold cross-validation procedure (iteratively training on 80% of trials and testing on the remaining 20%) and quantified the probability of each finger press being correctly identified within trials for each finger press sequence. The data was than averaged across sequences and z-scored across sensors or voxels within each participant. Sensor accuracy results were then plotted as t-statistic at the scalp level and significant sensors at p<0.01 marked with a point and a label. At the source level, whole brain results were corrected using a random effects analysis with an uncorrected threshold of t(15) > 3.73, p<0.001. Peak voxels and t-values of significant clusters were then reported in the results and the data plotted as t-statistics at the scalp level with a threshold of t(15) > 5.93, *p* < 0.05 (FWE corrected at the voxel level) centred on the peak voxel (MNI coordinates: −22, −38, 52) ^52^. The data thresholded at t(15) > 3.73, p<0.001 (uncorrected) can be found in Supplementary Figure 3b.

## Acknowledgements

We thank George Houghton, Ken Valyear, Simon Watt, Charlotte Stagg and Dario Farina for feedback on the findings. This study was supported by the Wellcome Trust Principle Research Fellowship to NB and the Sir Henry Wellcome Fellowship to KK.

## Author Contributions

K.K., D.B., G.B., N.B. designed the research plan. K.K. trained all participants. K.K., D.B.,S.M. performed the MEG recordings. K.K. and A.S. performed the EMG recordings. K.K. analysed the behavioural and EMG data. K.K and D.B. analysed the MEG surface data, D.B. and G.B. analysed the source data, K.K. prepared the figures. K.K., D.B., S.M., A.S., G.B. and N.B. wrote the manuscript.

---

**Supplementary Figure 1:**
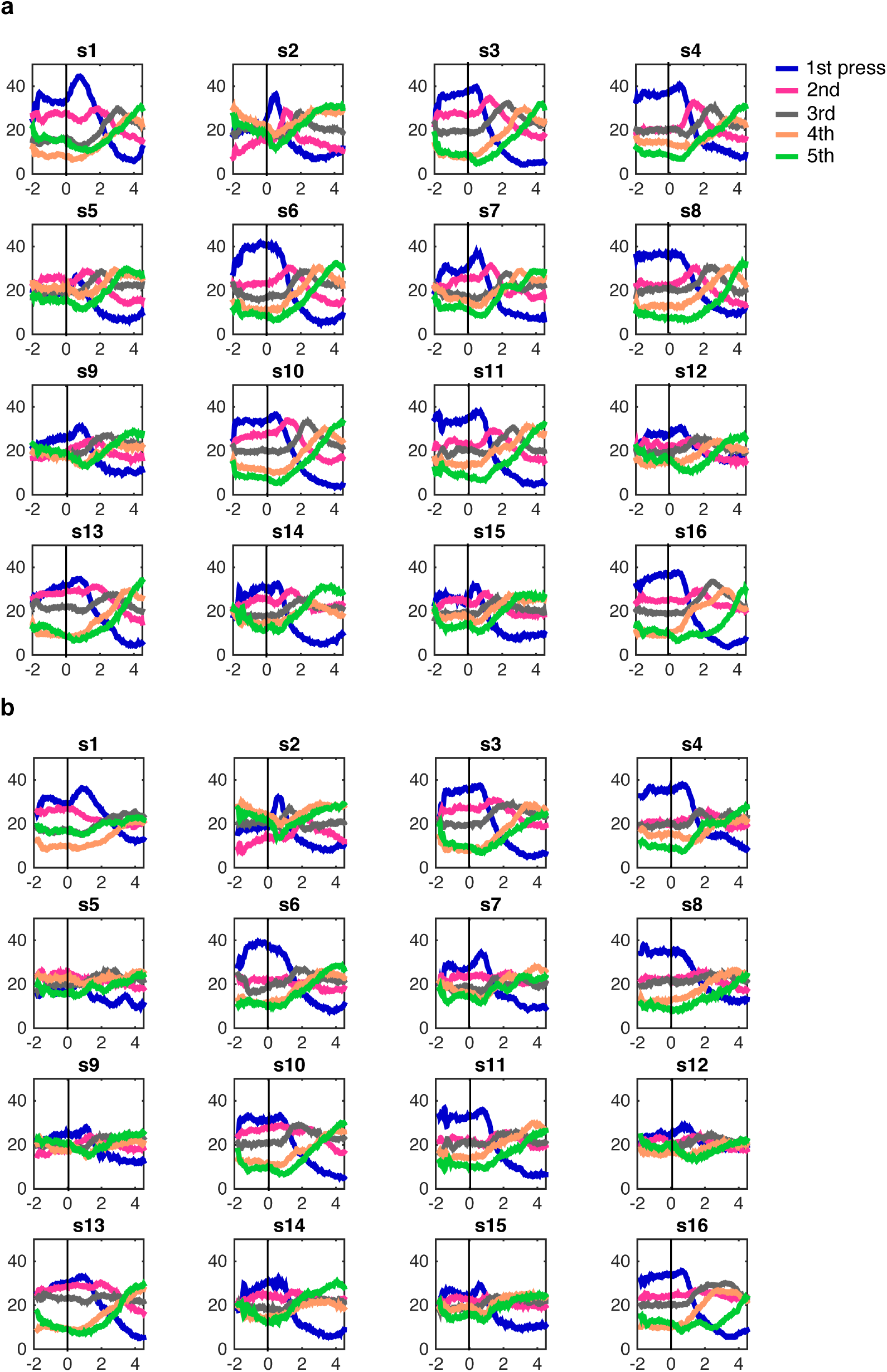
Individual press pattern probability traceplots. Trace plots for individual participants display the probability of each pattern classification related to the 1st-5th finger press 2000ms before and 4500ms after the ‘Go’ cue **a.** ‘within’ and **b.** ‘across’ finger sequences with the same target timing.

**Supplementary Figure 2:**
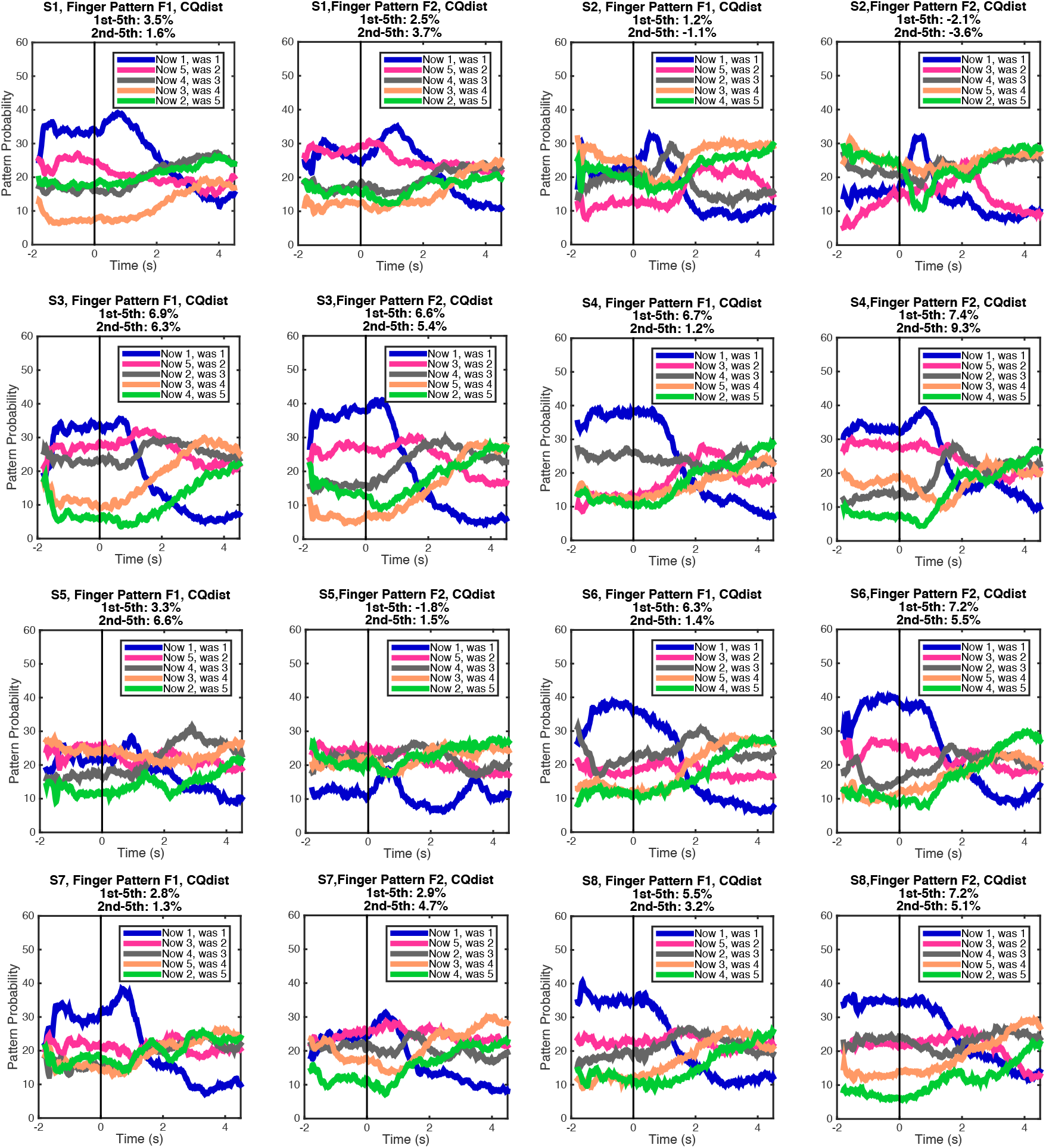
Effects of positional shifts of finger presses ‘across’ sequences with the same timing, but a different finger order. Individual traceplots show the ‘across’ sequence pattern probability labeled by position in the classified (‘now’) and trained (‘was’) sequence, respectively. In each participant, the graphs are split by finger order (F1 and F2), and indicate the temporal competitive queuing distance for each sequence in percent, respectively.

**Supplementary Figure 3:**
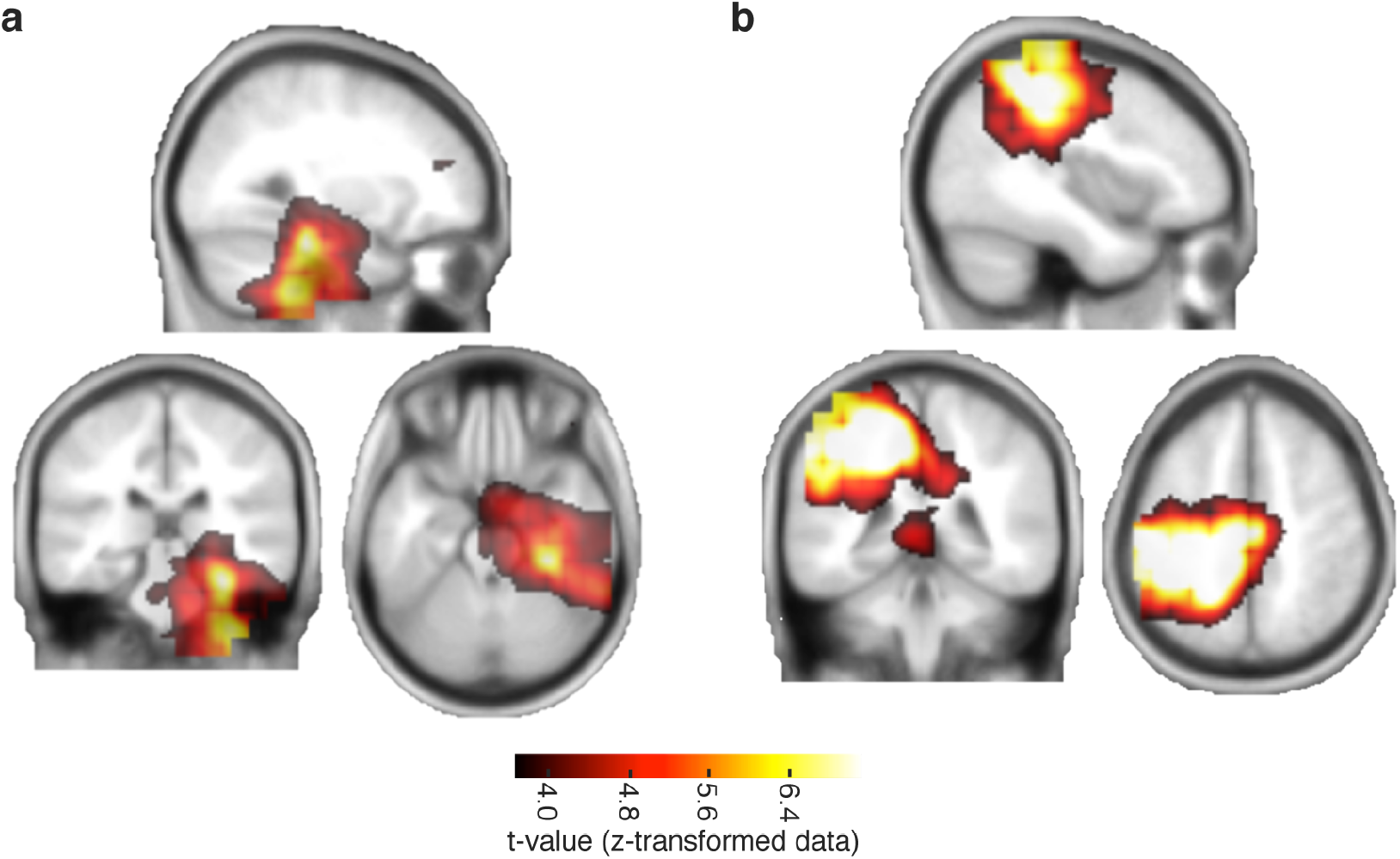
The source data as in Figure 7c-d plotted at a lower threshold of t(15) > 3.73, p <.001 (uncorrected) and centered at the peak voxel (MNI coordinates: 30, −30, −24) in panel **a.** and voxel coordinates: −22, −36, 52) in panel **b.**coordinates: −22, −36, 52) in panel **b.**

## References

1. Cermak, S. Developmental Dyspraxia. Adv. Psychol. (1985). doi:10.1016/S0166-4115(08)61143-7

2. Miller, N. The assessment of limb dyspraxia. Clin. Rehabil. (1988). doi:10.1177/026921558800200301

3. Stein, M. B., Baird, A. & Walker, J. R. Social phobia in adults with stuttering. Am. J. Psychiatry (1996). doi:10.1176/ajp.153.2.278

4. Sadnicka, A., Kornysheva, K., Rothwell, J. C. & Edwards, M. J. A unifying motor control framework for task-specific dystonia. Nature Reviews Neurology (2018). doi :10.1038/nrneurol.2017.146

5. Dressler, D. et al. Strategies for treatment of dystonia. Journal of Neural Transmission (2016). doi:10.1007/s00702-015-1453-x

6. Ebbinghaus, H. Memory: A Contribution to Experimental Psychology. Ann. Neurosci. (2013). doi:10.5214/ans.0972.7531.200408

7. Terrace, H. S. The simultaneous chain: A new approach to serial learning. Trends in Cognitive Sciences (2005). doi:10.1016/j.tics.2005.02.003

8. Sussillo, D., Churchland, M. M., Kaufman, M. T. & Shenoy, K. V. A neural network that finds a naturalistic solution for the production of muscle activity. Nat. Neurosci. 18, 1025–1033 (2015).

9. Shenoy, K. V, Sahani, M. & Churchland, M. M. Cortical control of arm movements: a dynamical systems perspective. Annu. Rev. Neurosci. 36, 337–59 (2013).

10. Laje, R. & Buonomano, D. V. Robust timing and motor patterns by taming chaos in recurrent neural networks. Nat. Neurosci. 16, 925–933 (2013).

11. Goudar, V. & Buonomano, D. V. Encoding sensory and motor patterns as time-invariant trajectories in recurrent neural networks. Elife 7, e31134 (2018).

12. Lashley, K. S. in Cerebral mechanisms in behavior 112–131 (Wiley, 1951).

13. Rhodes, B. J., Bullock, D., Verwey, W. B., Averbeck, B. B. & Page, M. P. A. Learning and production of movement sequences: Behavioral, neurophysiological, and modeling perspectives. Hum. Mov. Sci. 23, 699–746 (2004).

14. Page, M. P. A. & Norris, D. The Primacy Model: A New Model of Immediate Serial Recall. Psychol. Rev. (1998). doi:10.1037/0033-295X.105.4.761-781

15. Hartley, T. & Houghton, G. A linguistically constrained model of short-term memory for nonwords. J. Mem. Lang. 35, 1–31 (1996).

16. Burgess, N. & Hitch, G. J. Memory for serial order: A network model of the phonological loop and its timing. Psychol. Rev. 106, 551–581 (1999).

17. Botvinick, M. M. & Plaut, D. C. Short-term memory for serial order: A Recurrent Neural Network Model. Psychol. Rev. (2006). doi:10.1037/0033-295x.113.2.201

18. Averbeck, B. B., Chafee, M. V., Crowe, D. A. & Georgopoulos, A. P. Parallel processing of serial movements in prefrontal cortex. Proc. Natl. Acad. Sci. 99, 13172–13177 (2002).

19. Bohland, J. W., Bullock, D. & Guenther, F. H. Neural Representations and Mechanisms for the Performance of Simple Speech Sequences. J. Cogn. Neurosci. 22, 1504–1529 (2010).

20. Bullock, D. Adaptive neural models of queuing and timing in fluent action. Trends in Cognitive Sciences 8, 426–433 (2004).

21. Burgess, N. & Hitch, G. Computational models of working memory: Putting long-term memory into context. Trends Cogn. Sci. 9, 535–541 (2005).

22. Kornysheva, K. & Diedrichsen, J. Human premotor regions parse sequences into their spatial and temporal features. Elife 3:e03043, (2014).

23. Kornysheva, K., Sierk, A. & Diedrichsen, J. Interaction of temporal and ordinal representations in movement sequences. J. Neurophysiol. 109, 1416–24 (2013).

24. Ullén, F. & Bengtsson, S.L. Independent processing of the temporal and ordinal structure of movement sequences. J. Neurophysiol. 90, 3725–35 (2003).

25. Wiestler, T. & Diedrichsen, J. Skill learning strengthens cortical representations of motor sequences. Elife 2, e00801 (2013).

26. Wiestler, T., McGonigle, D. J. & Diedrichsen, J. Integration of sensory and motor representations of single fingers in the human cerebellum. J. Neurophysiol. 105, 3042–53 (2011).

27. Henson, R. N. . Short-term memory for serial order. Unpubl. Dr. Diss. Univ. Cambridge, Engl. 137, 73–137 (1996).

28. Binder, E. et al. Sensory-guided motor tasks benefit from mental training based on serial prediction. Neuropsychologia 54, 18–27 (2014).

29. Ali, F. et al. The basal ganglia is necessary for learning spectral, but not temporal, features of birdsong. Neuron 80, 494–506 (2013).

30. Diedrichsen, J., Criscimagna-Hemminger, S. E. & Shadmehr, R. Dissociating Timing and Coordination as Functions of the Cerebellum. J. Neurosci. 27, 6291–6301 (2007).

31. Zelaznik, H. N. et al. Timing variability in circle drawing and tapping: Probing the relationship between event and emergent timing. J. Mot. Behav. 37, 395–403 (2005).

32. Ivry, R. B., Spencer, R. M., Zelaznik, H. N. & Diedrichsen, J. The cerebellum and event timing. Ann. N. Y. Acad. Sci. 978, 302–317 (2002).

33. Conditt, M. a & Mussa-Ivaldi, F. a. Central representation of time during motor learning. Proc. Natl. Acad. Sci. U. S. A. 96, 11625–30 (1999).

34. Bullock, D. Adaptive neural models of queuing and timing in fluent action. Trends Cogn. Sci. 8, 426–433 (2004).

35. Hartley, T., Hurlstone, M. J. & Hitch, G. J. Effects of rhythm on memory for spoken sequences: A model and tests of its stimulus-driven mechanism. Cogn. Psychol. 87, 135–178 (2016).

36. Kornysheva, K. & Diedrichsen, J. Human premotor areas parse sequences into their spatial and temporal features. Elife 3, e03043 (2014).

37. Bengtsson, S. L., Ehrsson, H. H., Forssberg, H. & Ullén, F. Effector-independent voluntary timing: behavioural and neuroimaging evidence. Eur. J. Neurosci. 22, 3255–3265 (2005).

38. Buonomano, D. V. & Laje, R. Population Clocks: Motor Timing with Neural Dynamics. Space, Time Number Brain 14, 71–85 (2011).

39. Lungu, O. et al. Striatal and hippocampal involvement in motor sequence chunking depends on the learning strategy. PLoS One 9, e103885 (2014).

40. Steele, C. J. & Penhune, V. B. Specific increases within global decreases: a functional magnetic resonance imaging investigation of five days of motor sequence learning. J. Neurosci. 30, 8332–41 (2010).

41. Schendan, H. E., Searl, M. M., Melrose, R. J. & Stern, C. E. An FMRI study of the role of the medial temporal lobe in implicit and explicit sequence learning. Neuron (2003). doi:S0896627303001235 [pii]

42. Kraus, B. J., Robinson, R. J., White, J. A., Eichenbaum, H. & Hasselmo, M. E. Hippocampal ‘Time Cells’: Time versus Path Integration. Neuron (2013). doi:10.1016/j.neuron.2013.04.015

43. Jacobs, N. S., Allen, T. A., Nguyen, N. & Fortin, N. J. Critical role of the hippocampus in memory for elapsed time. J. Neurosci. (2013). doi:10.1523/JNEUROSCI.1733-13.2013

44. Friston, K. & Buzsáki, G. The Functional Anatomy of Time: What and When in the Brain. Trends Cogn. Sci. 20, 500–511 (2016).

45. Manns, J. R., Howard, M. W. & Eichenbaum, H. Gradual Changes in Hippocampal Activity Support Remembering the Order of Events. Neuron (2007). doi:10.1016/j.neuron.2007.08.017

46. De Zeeuw, C. I. et al. Spatiotemporal firing patterns in the cerebellum. Nat. Rev. Neurosci. 12, 327–44 (2011).

47. Gao, Z., van Beugen, B. J. & De Zeeuw, C. I. Distributed synergistic plasticity and cerebellar learning. Nat. Rev. Neurosci. 13, 619–35 (2012).

48. Hochberg, L. R. et al. Neuronal ensemble control of prosthetic devices by a human with tetraplegia. Nature (2006). doi:10.1038/nature04970

49. Shanechi, M. M. et al. Neural population partitioning and a concurrent brain-machine interface for sequential motor function. Nat. Neurosci. 15, 1715–1722 (2012).

50. Litvak, V. et al. EEG and MEG data analysis in SPM8. Comput. Intell. Neurosci. (2011). doi: 10.1155/2011/852961

51. Oostenveld, R., Fries, P., Maris, E. & Schoffelen, J. M. FieldTrip: Open source software for advanced analysis of MEG, EEG, and invasive electrophysiological data. Comput. Intell. Neurosci. (2011). doi:10.1155/2011/156869

52. Worsley, K. J. J. et al. A unified statistical approach for determining significant signals in images of cerebral activation. Hum. Brain Mapp. 4, 58–73 (1996).

